# Molecular circuit between *Aspergillus nidulans* transcription factors MsnA and VelB to coordinate fungal stress and developmental responses

**DOI:** 10.1101/2025.01.16.633343

**Authors:** Emmanouil Bastakis, Jennifer Gerke, Seyma Özkan, Rebekka Harting, Tanja Lienard, Christoph Sasse, Emmanouil S. Xylakis, Merle Aden, Anja Strohdiek, Gabriele Heinrich, Verena Grosse, Gerhard H. Braus

## Abstract

Development and secondary metabolism of the filamentous fungus *Aspergillus nidulans* are tightly controlled by concerted actions of several master regulator transcription factors. The connection between fungal development and cellular stress response programs is often elusive. Here we show that the MsnA zinc finger transcription factor, which controls salt-stress response, is a novel major player in fungal development. A molecular circuit among MsnA and the velvet domain regulator VelB was discovered, which mutually fosters the actions of both regulatory proteins during development. MsnA controls the expression of several genes encoding master transcriptional regulators of asexual as well as sexual development. In addition, MsnA affects directly and indirectly the synthesis of specific secondary metabolites relevant for fungal defense against other organisms and growth, in addition to salt-stress responses. Moreover, the expression of genes encoding the epigenetic regulators VapA and VipC are also directly controlled by MsnA. These subunits of the VapA-VipC-VapB methyltransferase signal transduction complex promote asexual differentiation. MsnA is therefore placed at a novel prominent position of the central regulatory network, which coordinates stress responses with the developmental and metabolic fate of the fungus.

## Introduction

Fungi are sessile forms of life, which have to react to various environmental signals in order to survive and propagate themselves successfully. Stress signals have to be perceived and mediated by several regulatory proteins to change gene expression and induce specific responses. At the end of these pathways are transcription factors (TFs), which control the gene expression for proteins, which are tightly associated with specific stress responses.

The velvet-domain master regulators form a group of TFs with roles in the development and the secondary metabolism of numerous fungi, particularly studied in *A. nidulans* [1]. These proteins can interact with DNA through their velvet-domain, which has the same structural fold as the Rel domain of the mammalian NF-κB transcription factor [2]. The four known velvets in *A. nidulans* are VeA (velvet protein A), VelB (velvet-like B), VosA (viability of spores A) and VelC (velvet-like C). The ability of the velvet proteins to form homo- or hetero-dimers have been identified as a necessary property of them. These velvet dimer formations define the shifting between the developmental programs and the regulation of secondary metabolite synthesis [1,3–5].

Apart from the velvet master regulators, the trimeric complex of VapA-VipC-VapB, plays an important role, specifically in the initiation of *A. nidulans* asexual development [6]. This complex is tethered to the zinc finger plasma membrane protein VapA. Upon external signals the methyltransferase heterodimer VipC-VapB is released from the membrane and can act subsequently in two ways. It interacts with VeA in the cytoplasm, hence, preventing it from entering the nucleus to induce the expression of genes promoting sexual development. VipC-VapB also enters the nucleus and via its methytransferase activity can decrease the H3K9me3 epigenetic mark in regulatory regions of genes encoding asexual master regulators such as BrlA (Bristle A) and AbaA (Abacus A). This initiates the asexual developmental program. The importance of the velvet proteins or the VapA-VipC-VapB trimeric complex in fungal development is known since years. However, the role of other potentially important but so far elusive developmental regulators and their prospective interactions with these already established networks remains to be discovered.

Msn2 and Msn4 are functionally redundant Cys2His2 (C2H2) zinc-finger TFs implicated into stress responses in *Saccharomyces cerevisiae* [7]. Their functions are related to the recognition of specific stress-response elements DNA elements (STREs), located to promoters of stress-related genes. The activation of the corresponding genes can be induced through this association and mediates a specific response to different stress stimuli at a time. The *A. nidulans* MsnA protein represents an orthologue of *S. cerevisiae* Msn2 or Msn4, which is also linked to stress responses, indicating a conserved function through the fungal kingdom [8]. Another study focusing on *A. parasiticus* and *A. flavus* showed an implication of MsnA in fungal development. In fact, *msnA* deletion strains showed an increased number of conidia compared to wildtype for both *Aspergilli* suggesting an inhibiting function on asexual development [9]. MsnA from *A. parasiticus* is further linked to the regulation of secondary metabolite genes associated with the synthesis of the carcinogenic mycotoxin aflatoxin as part of the fungal defense, and therefore a much broader response than only to oxidative stress [10]. Little is known regarding the roles in and the contribution of MsnA in the development of *A. nidulans*, whereas the relationship of MsnA and its paralogues to stress response has been extensively studied in various fungi. Any potential interplay of MsnA and known master regulators of *A. nidulans* development remains yet elusive.

This study revealed that *A. nidulans* MsnA is a *bona fide* transcriptional regulator that strongly binds in vivo directly to promoters of genes for asexual and sexual development and also fine tunes genes for the appropriate fungal secondary metabolism. MsnA directly controls transcription of the epigenetic VapA-VipC-VapB methytransferase complex and therefore promotes asexual fungal development. In summary, a novel genetic and molecular interplay among the MsnA and VelB fungal master regulators was discovered.

## Results

### The salt stress nuclear regulator MsnA is required for *A. nidulans* development

Developmental processes are tightly connected to various stress signals in filamentous fungi. NADPH oxidases generate reactive oxygen species (ROS), which induce fungal sexual development [11,12]. Cellular complexes as the COP9 signalosome connect protein stability control [13] to transcriptional and metabolic responses as well as to hormones and oxidative stress protection and ultimately developmental programs [14].

The Msn2 and Mns4 Cys2His2 (C2H2) zinc finger transcription factors (TFs) of the yeast *S. cerevisiae* or its counterpart MsnA in *A. nidulans* control various stress responses [7,8]. However, the molecular mechanisms linking stress response and developmental control or the contribution of MsnA (AN1652) to *A. nidulans* development are yet elusive. The *A. nidulans msnA* gene encodes an open reading frame of 1807 nucleotides interrupted by a single intron of 64 nucleotides (Fig 1A). The deducing 580 amino acids protein includes two Cys2His2 (C2H2)-type zinc finger DNA binding domains as was found in InterPro data base [15] , a nuclear localization signal (NLS), as predicted by the cNLS mapper tool [16], as well as a nuclear export signal (NES), as discovered by the LocNES algorithm [17] (Fig 1A). At least six phosphosites were predicted by the NetPhos tool [18] across the MsnA protein that might constitute direct kinase targets for phosphorylation. Additionally, there is at least one lysine residue predicted *in silico* by the UbiProber tool [19] representing a strong candidate site for a ubiquitination degradation signal recognized by the 26S proteasome.

**Fig 1.**
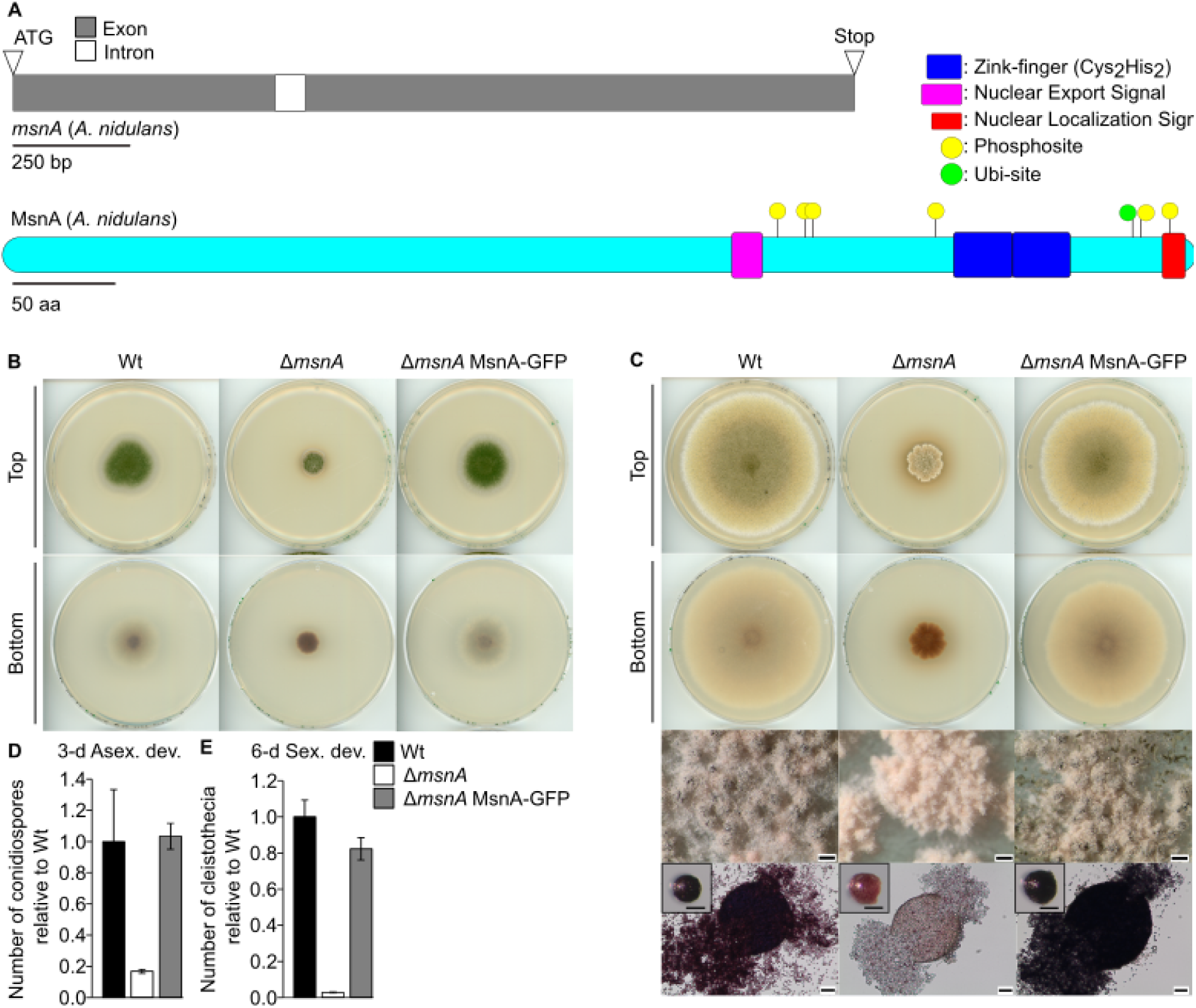
The MsnA stress response transcription factor is required for the development of *A. nidulans*. (**A**) Upper part open reading frame of *msnA* (AN1652), lower part protein domain analysis and predicted sites for post-translational modifications of MsnA. Blue rectangle = Zink-finger (Cys2His2) domain, purple rectangle = NES (Nuclear Export Signal), red rectangle = NLS (Nuclear Localized Signal), yellow circles = putative phosphosites and green circle = putative ubiquitination site. Phenotypical analysis of *A. nidulans* Δ*msnA* and complementation strain (Δ*msnA* MsnA-GFP), under (**B**) 3-d asexual (constant light) and (**C**) 6-d sexual (dark) promoting growth conditions. Scans derive from initial spots of 2000 conidia in the middle of the plate, which were then grow at 37 °C. Scale bars in overview photos (lower part of panel **C**) of many cleistothecia show a size of 200 µm, the bars shown in images of single intact cleistothecia represent 100 µm length, and the ones included in the images of broken cleistothecia represent 50 µm length. Quantification of (**D**) conidia and (**E**) cleistothecia of wildtype (Wt), Δ*msnA* mutant and complementation strain Δ*msnA* MsnA-GFP, after the indicative number of days, during asexual and sexual development. A total of 30,000 spores from each strain were spread equally (with glass beads) on each plate, and a minimum of three different plates were used per strain for the quantification.

An *msnA* deletion strain was generated to gain a better understanding of the contribution of MsnA to *A. nidulans* growth and differentiation programs (Fig 1B and 1C). The deletion strain was complemented with a functional MsnA-GFP (green fluorescent protein) fusion encoding gene driven by its native promoter (complementation strain), which resulted in wildtype-like development. In contrast, the Δ*msnA* strain was severely affected in colony growth (Fig 1B and 1C). The deletion strain showed an altered colony morphology at the outer borders, which appear to be uneven. Colony growth was additionally much slower compared to wildtype and complementation strain, which is also visible upon longer incubation of the plates for 8 days under asexual or 12 days of sexual development inducing conditions (S1A and 1B Fig). The *msnA* deletion strain additionally showed changes in colony color, suggesting an altered secondary metabolism. The number of conidia produced after 3 days of asexual development is severely decreased in the deletion strain when compared to wildtype and complementation strain (Fig 1D). After 6 days of sexual development, the Δ*msnA* strain additionally produced reduced numbers of the closed sexual fruiting bodies of the fungus. These cleistothecia, which are normally the overwintering structures in the soil appeared to be immature (Fig 1C and 1E). Incubation of the strains for 12 days, resulted in the formation of mature cleistothecia with sexual ascospores (S1B Fig), suggesting that resting structures production is delayed. These data support an important role for MsnA in *A. nidulans* growth and development.

Cellular localization of the functional MsnA-GFP fluorescence signal was determined in submerged cultures of the complementation strain after 20 hours growth in vegetative phase. The MsnA-GFP localization is nuclear, because it clearly colocalized with the nuclear dye of Hoechest already during early hyphal growth (S1C Fig). In summary, MsnA contributes to normal colony formation as well as sexual and asexual development.

### Genome-wide in vivo binding profiling of MsnA highlights its direct influence in various gene regulatory networks

The binding landscape of Msn2 from *S. cerevisiae*, particularly under stressful conditions has been studied in vivo in the past [20,21]. However, our current knowledge is restricted regarding the in vivo direct target genes of MsnA in *A. nidulans*. A ChIP-seq (chromatin immunoprecipitation coupled with next generation sequencing) experiment was performed by using the MsnA-GFP complementation strain with mycelia grown in submerged cultures for 20 hours at 37 °C. Statistically significant peaks (p<0.05 and fold enrichment (F.E.) ≥ 2.0) were identified as regions where there is significant alignment and enrichment of reads of the sample derived from the IPs with GFP-antibody of MsnA-GFP compared to the input control samples where no antibody was applied. The peaks were located to regions spanning up to 3 kb upstream from the transcription start sites (TSS) of 1439 distinct unique locus IDs (genes) in two independent sets of ChIP-seq analysis, consisting each of the MsnA-GFP versus its input sample (Fig 2A and S4 Table). It was further examined, how the peaks, identified from the ChIP-seq, are distributed over different genetic elements of the genome. Around 85 % of the identified peaks were found to be located to promoter regions of genes up to 3 kb from the TSS. The great majority of the binding events were mostly located in promoter regions spanning at around 1 kb upstream from the nearby genes (Fig 2B).

**Fig 2.**
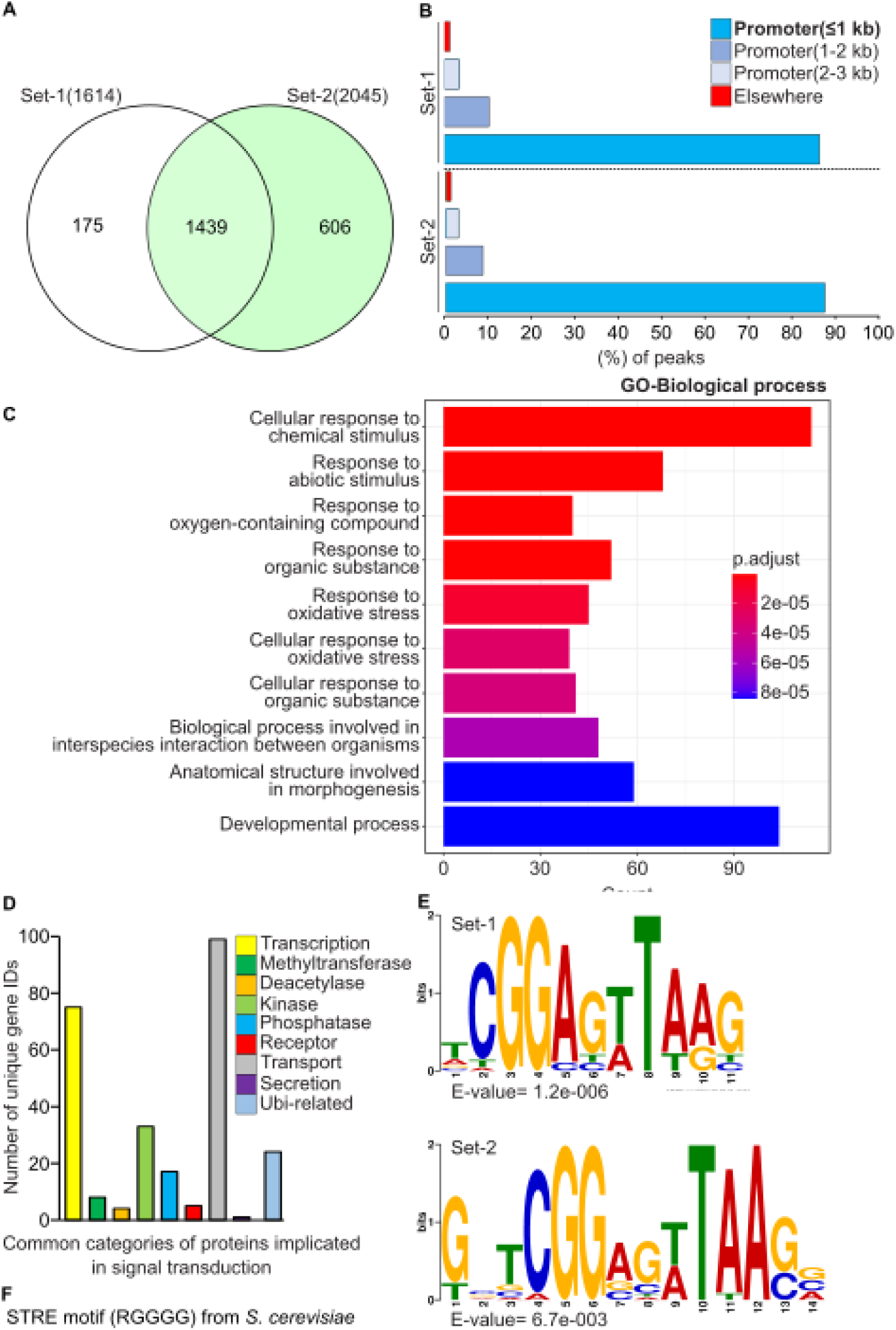
Genome-wide in vivo binding profile of MsnA-GFP in *A. nidulans*. (**A**) The Venn diagram depicts the overlap of two independent sets of ChIP-seq analysis (MsnA-GFP versus input). A total number of 1439 locus IDs of genes (cut offs: fold enrichment (F.E.) ≥ 2.0, and with p < 0.05), associated with peaks (located in up to 3 kb promoter regions) found simultaneously in both independent sets of the ChIP-seq samples. MsnA-GFP was detected by the GFP antibody and the corresponding input samples where used as negative control. (**B**) Bar charts show for both independent sets of the ChIP-seq analyses that more than 85 % of the statistically significant identified peaks are falling into promoter regions of genes spanning up to 1 kb from the TSS (transcription start site). (**C**) Bar plots show the GO (Gene Ontology)-enrichment analysis, in terms of the biological processes for the 1439 common IDs of genes identified simultaneously in both sets of the analyses. (**D**) Bar diagram shows genes encoding proteins of signal transduction processes. The numbers correspond to unique genes found after manual search in the set of the 1439 common IDs identified from both sets of analyses. (**E**) Logos depict the top ranked *de novo* motifs, discovered by using the RSAT-peak-motif tool. For each independent set a group of the 100 bp sequences was used as input of the RSAT tool. Each of these sequences were located underneath the summit of the top 150 ChIP-seq peaks. All of those peaks were located into 3 kb promoter regions. (**F**) the STRE (stress response element) DNA recognition motif of Msn2 from *S. cerevisiae*.

The different types of proteins encoded by the genes, which are presumably targeted by MsnA, were determined and classified. A GO (Gene Ontology)-enrichment analysis was performed with all genes found to be associated to MsnA within a range of 3 kb upstream from the corresponding transcription start site. The GO analysis showed that the most prominent and highly statistically significant target genes are coding for fungal proteins implicated in biological processes (BP) as development, morphogenesis and cellular responses towards chemical or abiotic stimuli, respectively (Fig 2C). GO analysis with a focus on the cellular component mode of the MsnA target proteins revealed as most representative categories the cell periphery, the plasma membrane and the extracellular region of the fungus (S2 Fig). Search among nine protein categories, known to be associated with regulation or signal transduction, revealed top candidates related to transcription, phosphorylation or transport (Fig 2D). This corroborates a strong potential MsnA impact on the control of genes encoding proteins with direct implications to cellular signaling cascades.

The MsnA-associated DNA motifs found in *A. nidulans* were compared for similarities to the previously described stress response element (STRE) sequences from *in vitro* [7] or in vivo ([20] studies in *S. cerevisiae* (RGGGG motif). The RSAT-peak-motifs [22] webtool was employed. A *de novo* motif discovery was performed by using the 100 bp sequences laying under the peak’s summits of the top 150 identified peaks (based to their fold enrichment). The discovered DNA motif from *A. nidulans* was found to be very similar to the described yeast motif and is therefore conserved between yeasts and filamentous ascomycetes (Fig 2E and 2F).

In summary, these results from the ChIP-seq show that *A. nidulans* MsnA can recognize and subsequently be associated with STRE DNA motifs and can influence the expression of genes that are associated with signal transduction pathways.

### *A. nidulans* MsnA directly controls expression of genes encoding master regulators of fungal development

A fundamental step for the initiation of the asexual developmental program, which will lead to the formation and maturation of the asexual spores (conidia) is the activation of the *brlA* (*bristle A*) promoter for a master initial transcription factor for conidiation. This activation requires the derepression caused by the SfgA (suppressor of *fluG*) against the conidiation and the derepression that is caused by the occupation of the *brlA* promoter by the NsdD (never in sexual development D) and VosA (viability of spores A) transcription factors [23–25]. FluG (fluffy G) plays a major role for removing those repressive effects from the *brlA* promoter [26], hence, allowing the Flb (fluffy low brlA) regulators (FlbB, C, D and E) to activate directly the corresponding promoter [27–31]. FluG, SfgA and the Flbs are operating as upstream developmental activators (UDAs) for the *blrA* promoter [25]. Once the BrlA transcription factor is expressed, it controls (directly and indirectly), a development cascade of genes, known as central developmental pathway (CDP) [25]. BrlA directly induces *abaA* (*abacus A*) expression encoding the regulator responsible for the middle phase of sporulation [30]. AbaA in turn activates, among other genes, *wetA* (*wet-white A*), that effects the expression of genes with roles at late stages of conidiation [30,32–34]. The *stuA (stunted A),* a helix-loop-helix transcription factor and *rgsA* (*regulator of G protein signaling*) genes are two additional components of the CDP. The expression of *stuA* is BrlA-dependent and a deletion strain of *stuA* shows defective formation of conidiophores with only a few conidia. The RGS (regulator of G-protein signaling) proteins FlbA and RgsA are further promoting asexual development. In particular, FlbA alongside with FluG and the rest of the Flbs are required for the activation of *brlA* [35] and at the same time suppressing FadA, an inhibitor of asexual development [36]. RgsA is promoting conidia formation, by attenuating the suppressive effects that GanB shows against the asexual growth [37].

The *A. nidulans* sexual developmental program is tightly controlled by several regulators to produce the sexual reproductive spores (ascospores) in cleistothecia as enclosed fruiting bodies. The FadA (fluffy autolytic dominant A) protein, a member of the heterotrimeric G-protein (SfaD-GpgA-FadA) mediates signals from the plasma membrane to proteins (effectors) inside the cell [23]. When FadA is bound to GTP, it can initiate an MPA-kinase singling that leads to the activation of transcriptional master regulators of the sexual development like NsdD and SteA [38]. NsdD (GATA-like transcription factors), NsdC (C2H2-type transcription factor) and SteA (homeodomain-C2H2-zinc finger transcription factor) are three major regulators of the sexual development [39–41]. Deletion of each of these genes results in no cleistothecia formation, highlighting their essential role during the sexual development. Two additional transcriptional regulators with important roles during the sexual development are the Zn(II)2Cys6 transcription factors NosA (number of sexual spores) and RosA (repressor of sexual development) [42,43]. These two transcription factors have opposite roles, with NosA being an activator and RosA being a repressor of sexual development.

The MsnA-dependent developmental genetic network was approached by searching for in vivo binding events of MsnA to promoters of genes regulating asexual or sexual development as well. Binding of MsnA in vivo does not necessarily mean direct gene regulation. Therefore, comparative gene expression analysis by qRT-PCRs were performed between Δ*msnA* and wildtype to verify, which bindings correlate to changes in gene expression. Mycelia derived from submerged liquid cultures were analyzed using the same conditions as for the ChIP-seq experiments. Binding of MsnA to the promoters of all regulatory genes tested, except *fadA,* correlated with significant changes in transcripts expression after 20 hours of vegetative growth. These findings corroborate that the MsnA transcription factor is an early regulator in vegetative growth at the stage of developmental competence (Figs 3 and S3). MsnA rather acts as repressor than as inducer of these identified genes after 20 hours of vegetative growth, because 10 of in total 12 differentially expressed genes showed increased expression in the Δ*msnA* strain compared to only two genes with reduced transcription (Fig 3B and 3D).

**Fig 3.**
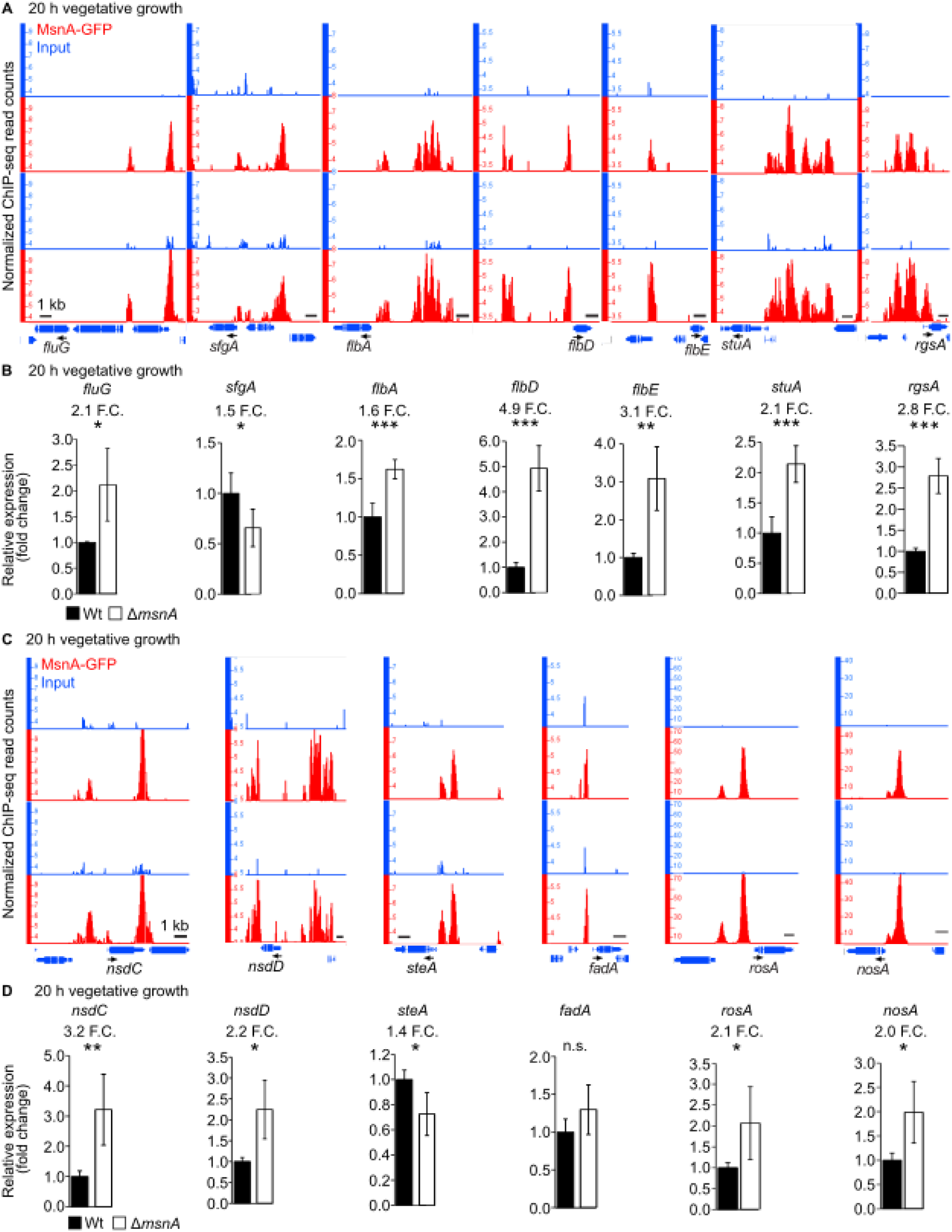
MsnA is associated in vivo with promoters and controls expression of genes encoding master regulators of *A. nidulans* asexual and sexual development, already during vegetative growth. Screen shots (**A** and **C**) from the Integrative Genome Browser (IGB) show peaks from the ChIP-seq data with MsnA-GFP located to the indicated promoters of regulatory genes. Small black horizontal arrows show the transcription direction of the corresponding gene below each screen shot. Further qRT-PCRs gene expression analyses (**B** and **D**) were performed, with RNA derived from wildtype (Wt) or Δ*msnA* strains grown for 20 hours vegetatively (same conditions with the ChIP-seq experiment). At least three biological qRT-PCR replicates were performed per strain and per different time point, respectively. Each biological replicate includes a minimum of three technical replicates. Statistical differences for the gene expression data were performed by t-test, with *: p<0.05, **: p<0.01 and ***: p<0.001, n.s.: not significant.

The impact of MsnA on developmental regulatory genes was further dissected by analysing early stages of asexual or sexual development. Gene expression was examined by qRT-PCRs with RNAs derived from wildtype (Wt) or Δ*msnA* strains, that were initially growing vegetatively and then shifted to solid medium plates to promote either the initiation of asexual or sexual development (S3A and S3B Fig). In most cases, the same genes as before were found to be differentially expressed at different time points by MsnA. An exception was the *rosA* expression during sexual growth (S3B Fig), which is not controlled by MsnA during sexual differentiation, but only during the vegetative phase (Fig 3D). Moreover, the expression of *fadA*, found not to be affected in mycelia growing vegetatively (Fig 3D), was also not affected during the swift of mycelia to sexual conditions for the indicative time points tested (S3B Fig).

SclB (sclerotia-like B) is a zinc cluster (C6) transcription factor with an important role in asexual development, secondary metabolism and oxidative stress [44]. SclB is characterized as an activator of the asexual developmental program, controlling the expression of *brlA* and several other genes encoding of regulatory proteins such as FluG and the FlbC and FlbD, operating upstream from BrlA [44].

SclB activates genes, which were also found to be direct targets of MsnA, such as *flbD* and *fluG* (Fig 3A to 3B). Therefore, MsnA might also be able to control the expression of *sclB* and its regulatory partner *brlA*. The search for peaks located proximal to *sclB* and *brlA* in the MnsA ChIP-seq data revealed a small, but clear peak in the promoter of *sclB* in close proximity with its TSS. No peak was detected close to the *brlA* proximal promoter region (S3C and S3F Fig). Examination of far away (tenths of kbs) upstream regions from the TSS of the promoters of both genes (*sclB* and *brlA*) revealed strong peaks indicating direct in vivo binding of MsnA. It cannot be excluded that binding to these remote regions can even contribute to the regulation of the corresponding downstream genes (S3C and S3F Fig).

Expression of *brlA* and *sclB* was investigated at the same conditions as for the ChIP-seq with MsnA. Expression of both genes was severely decreased in the Δ*msnA* strain after 20 hours vegetative growth (12.8 F.C. for the *sclB* and 3.8 F.C. for the *brlA*) compared to the wildtype strain (S3D and S3G Fig). In addition, the expression of *sclB* as well as of both functional overlapping *brlA* transcripts, *brlAα* and *brlAβ*, was examined at early stages of asexual development. Expression of *brlA* (including the two transcripts *brlAα* and *brlAβ*) and *sclB* genes was highly decreased at all tested time points during early asexual development in Δ*msnA* compared to wildtype strain (S3E and S3H Fig). These findings support a critical role of MsnA to increase gene expression for *brlA* and *sclB* to support the asexual developmental program.

### MsnA directly and indirectly controls fungal secondary metabolism

Secondary metabolism constitutes an essential part of fungal survival as chemical language to communicate with the environment [1,45] and is directly linked to development on a molecular level [4]. This includes the synthesis of chemical compounds that can be used as molecules with detrimental and often lethal effects against putative competitors or enemies of the fungus. Other bioactive molecules can either trigger or supress certain processes tightly associated with the developmental programs of the fungus [46,47].

The overall impact of MsnA as stress as well as developmental regulator on *A. nidulans* secondary metabolism was examined. Secondary metabolites from wildtype (Wt) and Δ*msnA* strains growing under asexual conditions for 2 days were extracted and resulting secondary metabolite profiles analyzed by LC-MS/MS were compared.

Austinol and dehydroaustinol were severely reduced in the absence of MsnA compared to wildtype. Especially dehydroaustinol promotes fungal asexual development [48] (Fig 4A). Moreover, the antimicrobial DHMBA (2,4-dihydroxy-3-methyl-6-(2-oxopropyl) benzaldehyde [49] was approximately eleven-fold increased in the *msnA* deletion strain compared to wildtype during the same growth conditions (Fig 4A). In conclusion, these data support multiple control functions of MsnA to adjust defense and signal secondary metabolite levels for supporting asexual fungal development. Secondary metabolites profiling of *ΔmsnA* versus the wildtype strain was examined additionally during vegetative and sexual growth of the fungus, but no significant differences were observed for any known metabolites.

**Fig 4.**
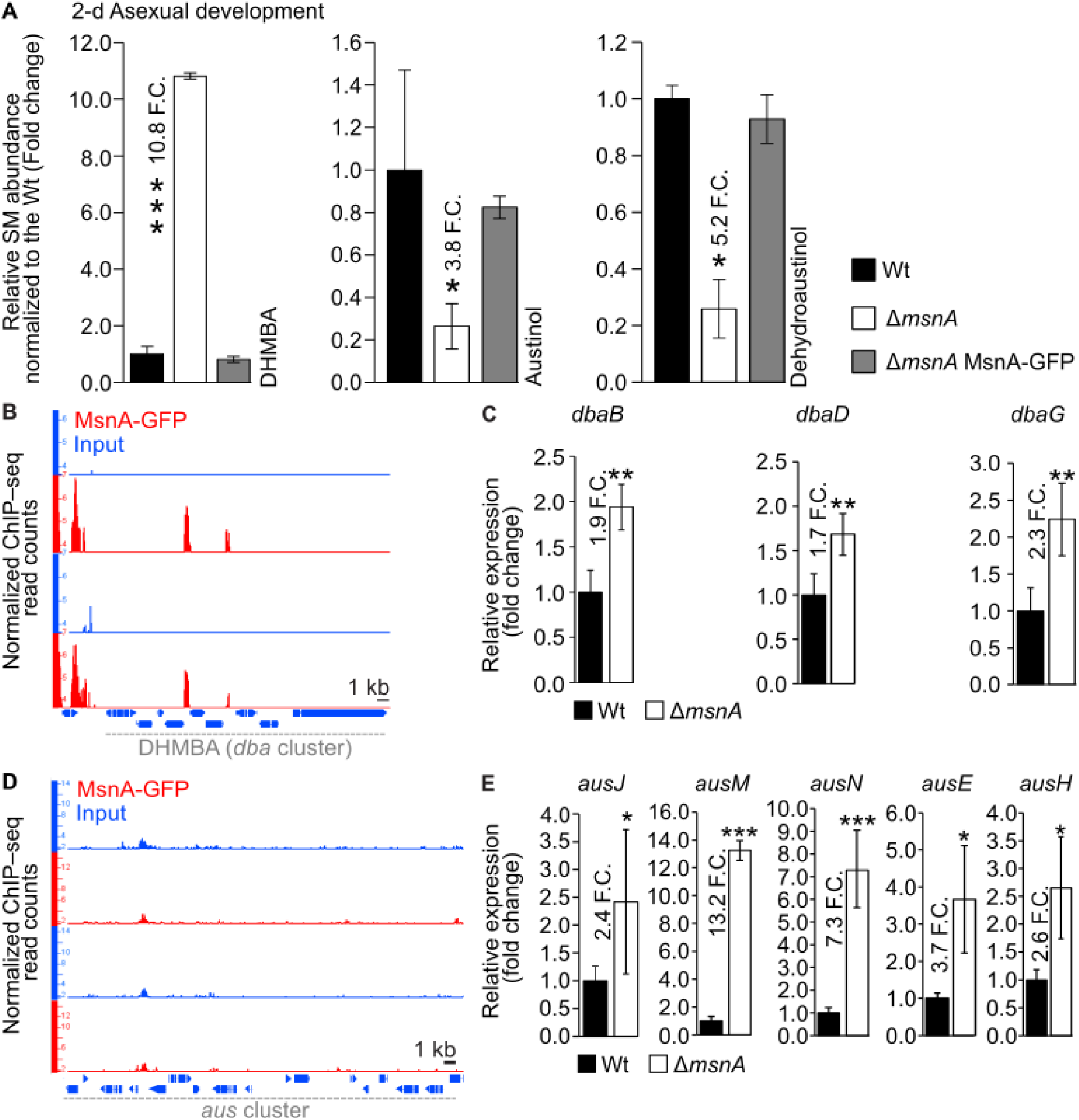
Fungal secondary metabolites associated with either defense or development of *A. nidulans* are controlled directly or indirectly by the stress regulator MsnA. Bar diagrams (**A**) depicting the relative abundance of secondary metabolites normalized towards wildtype; each strain is represented as the average and standard deviation of three replicates. Secondary metabolites were detected by LC-MS/MS with a charged aerosol detector (CAD) from extracts derived from the wildtype and the *msnA* deletion strain growing asexually for 2-d on minimal medium plates at 37°C. Screen shots from the Integrative Genome Browser (IGB) showing peaks from the ChIP-seq data with MsnA-GFP located at the promoters of genes of the *dba* (**B**)*, aus* (**D**) gene clusters. Gene expression was analyzed by qRT-PCR, for genes of the *dba* (**C**) and *aus* (**E**) gene clusters. All RNAs (for **C** and **E**) derived from mycelia of wildtype or Δ*msnA* strain, grown vegetatively in submerged cultures for 20 hours at 37° C and shaking by rotary shaker. Presented data represent average values and standard deviations of at least three biological replicates with each deriving from at least three technical replicates; Statistics made by Student’s t test: *P ≤ 0.05, **P ≤ 0.01 and ***P ≤ 0.01, n.s.: not significant.

The secondary metabolite profiling of the Δ*msnA* mutant strain revealed an important direct or indirect MsnA control function of the synthesis of enzymes of several known secondary metabolites. The ChIP-seq data were then searched for peaks that are potentially located to the promoters of the corresponding secondary metabolite gene clusters resulting in differentially synthesized molecules. Peaks were identified among the cluster genes that are associated with the formation of DHMBA, but not austinol and dehydroaustinol (Fig 4B and 4D).

Subsequently, it was investigated if peaks from the ChIP-seq data, found to be associated with secondary metabolite genes, are able to directly control the expression of genes for the synthesis of DHMBA, austinol and dehydroaustinol. Gene expression of wildtype and the Δ*msnA* strains was compared during vegetative growth. The two ChIP-seq peaks found inside the *dba* gene cluster (Fig 4B) support the hypothesis that MsnA directly controls transcriptionally the expression of several genes of the DHMBA gene cluster. A gene expression study confirmed that out of the nine in total genes that constitute the *dba* cluster, at least three showed significant changes in their gene expression in the Δ*msnA* strain compared to wildtype (Fig 4C). These changes in those genes expression indicate a clear repressive role of MsnA, which is in line with the increase of the DHMBA synthesis in the Δ*msnA* strain (Fig 4A).

Our study has also shown that there is a strong involvement of MsnA to the synthesis of specific metabolites such as austinol/dehydroaustinol (Fig 4A). However, no ChIP-seq peaks were found that indicate direct in vivo binding of MsnA to promoters of genes belonging to the corresponding secondary metabolite cluster (Fig 4D). Thus, there might be an indirect transcriptional regulation of this cluster by MsnA. Therefore, we examined the expression of genes that belong to the *aus* cluster when the fungus was growing vegetatively. The expression of the specific genes (*ausJ*, *M*, *N*, *E* and *H*) from the *aus* cluster, where their deletion leads to abolishment of austinol and dehydroaustinol synthesis [50], was found to be strongly increased in the Δ*msnA* strain (Fig 4E). These results confirmed an indirect transcriptional regulation of MsnA towards essential genes of the *aus* cluster, which is presumably mediated by other TFs downstream of MsnA. In conclusion, MsnA controls secondary metabolite cluster expression and subsequently secondary metabolites associated with fungal development and defense.

### An uncharacterized transcriptional circuit among the MsnA and the velvet master regulators

The impact of MsnA on *A. nidulans* development has to be embedded into the corresponding fungal regulatory networks including other known master regulators of differentiation as well as the response elements to external signals. Velvet domain proteins share a similar fold for DNA binding and dimerization which is reminiscent to mammalian NF-kB and are key regulators of fungal development [51]. The four velvet-domain proteins VeA, VelB, VelC and VosA can form various homo-and heterodimers with distinct function in fungal development. VelB can form a heterodimer with VeA [4], but can also associate to VosA [3].

The initial approach was to examine whether MsnA directly controls expression of any of the velvet genes. The MsnA-GFP ChIP-seq data were examined for peaks on the promoters of the velvet genes. Strong peaks were discovered in the promoters of *veA* and *velB*, but not in the promoters of *vosA* and *velC* (Fig 5A). This indicates a preference of MsnA to directly associate with regulatory regions of the *veA* and *velB* genes in vivo, where the encoded proteins can form the VelA-VelB heterodimer.

**Fig 5.**
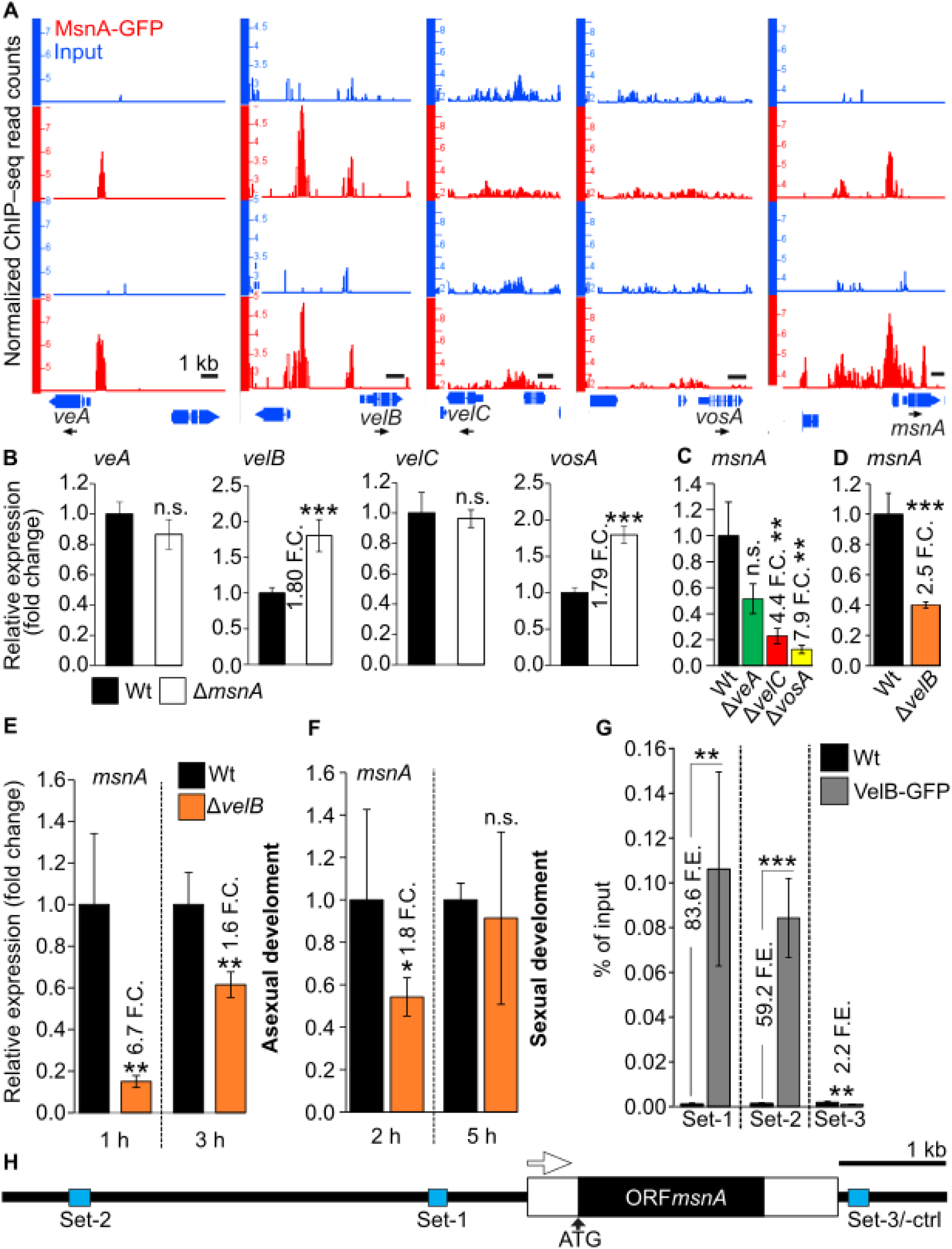
Stress regulator MsnA and velvet domain developmental regulators can reciprocally transcriptionally control each other. Screen shots from the Integrative Genome Browser (IGB) (**A**) showing peaks (binding events of MsnA) from the ChIP-seq data with MsnA-GFP associated with promoters of *veA* and *velB* genes. Gene expression analyses (**B**-**D**) of the *veA*, *velB*, *velC* and *vosA* and *msnA* via qRT-PCRs, with mycelia derived from submerged cultures of wildtype and the deletion strains Δ*msnA*, Δ*veA*, Δ*velC*, Δ*vosA*, and Δ*velB* grown vegetatively for 20 hours at 37° C shacking. Gene expression analyses of *msnA* from mycelia of wildtype (Wt) and Δ*velB*, grown initially for 20 hours under vegetative conditions and subsequently transferred to minimal medium plates for the indicated periods of time under (**E**) asexual (37 °C, light) and (**F**) sexual (37°C, dark, plates sealed with parafilm) conditions. ChIP-qRT-PCRs (**G**) for examining the in vivo binding of the VelB-GFP (expressed under its native promoter) to the promoter of *msnA*. The cartoon of the panel (**H**) depicts with blue boxes the different regions of DNA where the binding of VelB-GFP was tested by using distinct sets of primers at a case; set-1 and set-2 constitute regions where VelB-GFP is highly associated with, and set-3 is a region downstream of the *msnA* that was used as a negative control. For all the gene expression results and the ChIP-qRT-PCRs, presented data are averages and standard deviations of at least three biological replicates with each deriving from at least three technical replicates; Statistics made by Student’s t test: *P ≤ 0.05, **P ≤ 0.01 and ***P ≤ 0.01, n.s.: not significant.

A gene expression analysis was performed with mycelia grown for 20 hours vegetatively. It was examined whether presence or absence of MsnA affects gene expression of velvet domain protein transcripts. MsnA primarily reduces transcript formation of the *velB* as well as the *vosA* gene (Fig 5B). The in vivo binding of MsnA to the *veA* promoter, does not seem to induce differential expression of this gene, at least not at this particular time point (Fig 5A and 5B). Moreover, since there is no peak to the promoter of *vosA*, but there is a change in its expression in the Δ*msnA* compared to wildtype, it can be concluded that there is an indirect control of *vosA* by MsnA. Remarkably, the *velB* gene represents the only velvet gene where the MsnA is directly associated with its promoter resulting in differential gene expression (Fig 5A and 5B).

A comparative gene expression analysis of RNA, derived from mycelia growing for 20 hours vegetatively, was set to examine if velvets can affect the expression of *msnA*. This analysis revealed that the deletion strains of *veA*, *velC* and *vosA* regulators, showed a severely reduced expression of the *msnA* gene with the most prominent impact for the *vosA* deletion strain (Fig 5C). This suggests that VeA, VelC or VosA are required for sufficient amounts of *msnA* transcripts by either a direct or indirect molecular mechanism. A potential regulatory circuit between VelB and MsnA was examined in more detail, because the expression of the *velB* transcript is directly controlled by MsnA. Gene expression analysis with the wildtype and the *velB* deletion strain, was not only performed with mycelia that were grown under vegetative conditions but also with mycelia that were transferred onto solid medium after vegetative growth, which were then incubated under conditions that could favor either asexual or sexual development. Expression of *msnA* was found to be significantly repressed in the *velB* deletion strain (Δ*velB*) (Fig 5D). These results revealed that VelB was able to induce the expression of *msnA* during vegetative fungal growth, which supports a mutual control between both transcription factors.

VelB continues promoting the expression of *msnA* transcripts after the shift from vegetative growth to illuminated solid surface growth promoting asexual development for up to three hours with a peak after one hour (Fig 5E). The influence of MsnA to the expression of *velB* during sexual development inducing conditions in the dark on solid media is present at the outset of this developmental program, and five hours after it has been completely diminished (Fig 5F).

Lastly, since we discovered the VelB regulatory function for *msnA* expression, it was further examined whether this is an indirect control or based on a direct VelB in vivo binding to the *msnA* promoter region. A ChIP experiment was performed with mycelia from the VelB-GFP and the wildtype strain grown vegetatively for 20 hours. This revealed two *msnA* promoter positions (Fig 5G and 5H, set-1 and set-2) proximal to the TSS with strong in vivo VelB association in comparison to the control ChIP-qRT-PCR to a downstream region from *msnA*, where the ChIPed DNA of the IP with VelB-GFP was lower than the corresponding IP with wildtype (Fig 5G and 5H, set-3).

In summary, a novel mutually controlled genetic network between two key regulators of stress response (MsnA) and fungal development (VelB) was discovered. MsnA is also strongly associated in vivo with its own promoter (Fig 5A), which supports an additional level of autoregulation to further influence its own transcription. VelB and MsnA do not only control their own expression but also the expression of other key control genes for distinct developmental fungal programs through this network.

### *msnA* and *velB* genetically interact during fungal development

Genetic interactions between the *msnA* and *velB* genes were further explored due to the discovered molecular regulatory interplay among MsnA and VelB. The double deletion strain of *msnA* and *velB* was generated. Under asexual growth conditions the single mutants Δ*msnA* and Δ*velB* showed distinctively different phenotypes, regarding conidiospores formation and secondary metabolism (as can be observed macroscopically from the bottom of the plates) (Fig 6A). The phenotype of the Δ*velB* Δ*msnA* double mutant strain stands out, because it is a rather synergistic phenotype of both genes. More specifically, the examination of the *velB msnA* double deletion strain under conditions that favor the sexual development of the fungus showed prominent and distinctive phenotypes compared to the corresponding single mutants (Fig 6B). The *msnA* mutation has an epistatic role over *velB* regarding colony morphology (Fig 6B). In contrast, the role between *msnA* and *velB* is changing when we examined macroscopically their genetic interaction under the prism of secondary metabolism (coloration at the bottom of the colony), where *velB* acts epistatically towards *msnA* (Fig 6B). In summary, MsnA and VelB do not only mutually control their transcriptional regulation (Fig 5), but further share a variety of genetic interactions when growing under either asexual or sexual conditions.

**Fig 6.**
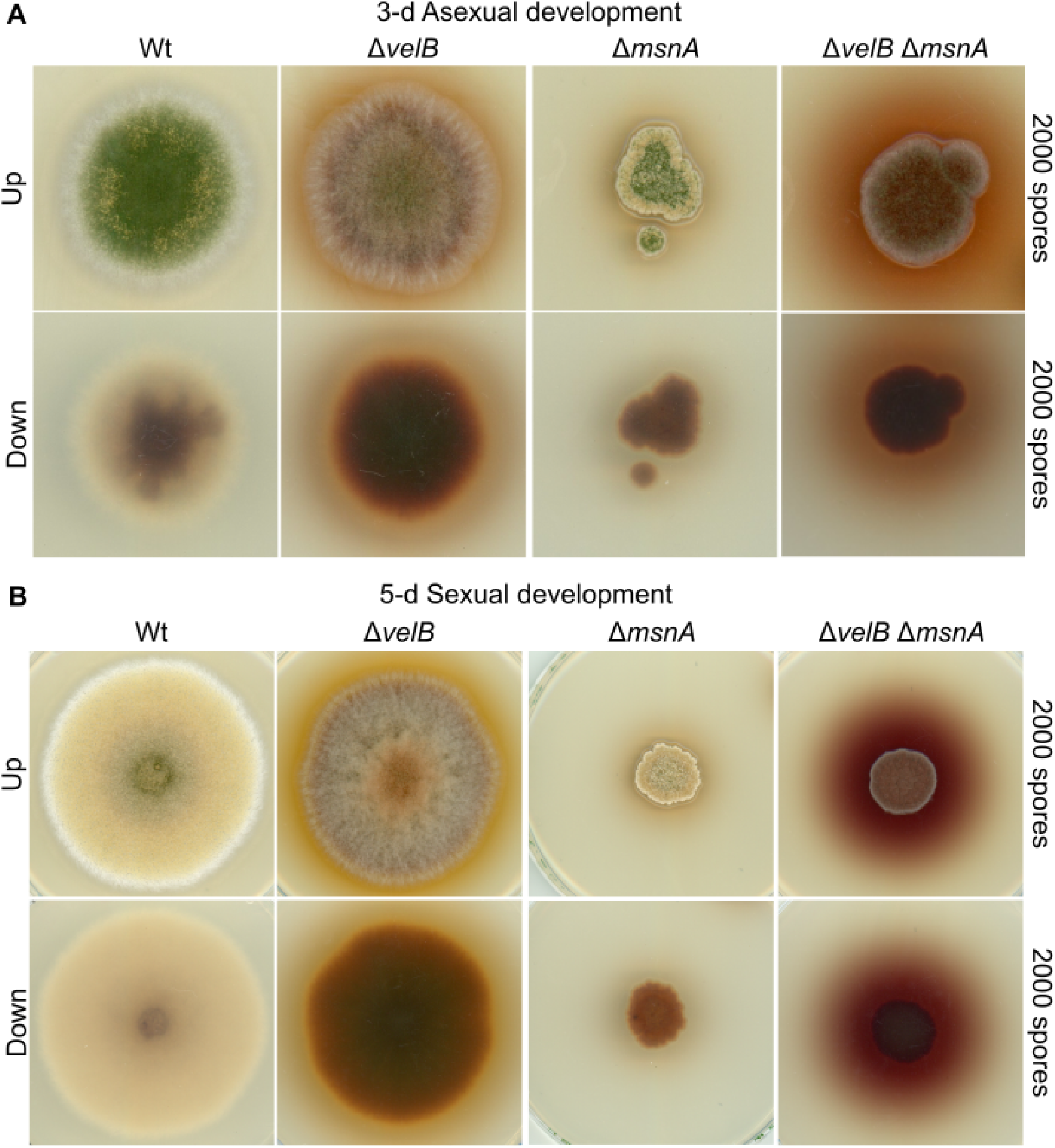
*msnA* genetically interacts with the *velB*. Scans of different strains grown under (**A**) asexual and (**B**) sexual conditions represent the phenotypes of the single and the corresponding double mutants of *msnA* and *velB*. For both phenotypical assays, spores from the depicted strains were initially spotted (2000 spores/plate) on minimal medium plates. Subsequently scans made from top of the plates after the indicated period of time, under either asexual or sexual growth conditions. Scans from the bottom of the plates highlight differences in the overall colorations of the fungal colonies, which is an indicator for changes secondary metabolism of the corresponding strains.

### MsnA controlled salt-stress response is independent of VelB

MsnA constitutes a central player in the salt-stress response of *A. nidulans* [8]. Based on our previous findings regarding the interaction of MsnA with VelB on a cross-regulatory and genetic level, it was hypothesized that such interactions might also play a role when the fungus is growing under salt-stress conditions. Therefore, the potential mutual control of VelB and MsnA was analyzed under salt stress. Single or double deletion strains of *msnA* and *velB* were cultivated upon solid medium plates containing 1 M NaCl under conditions that either induce asexual or sexual development. Diameters of stressed and non-stressed colonies of all the strains were measured and the ratios between mock and treatment were calculated. The salt treatment resulted in a retardation in colony growth for all strains (Fig 7A and 7B). The quantification of this effect showed that during asexual and sexual development they were striking differences, compared to wildtype (Wt), only for the strains that carry the Δ*msnA* (Fig 7C and 7D). In fact, the presence of the *velB* deletion did not contribute to the salt-stress response of the fungus, even when combined with the Δ*msnA*. Under the given stress conditions, during asexual and the sexual growth, *msnA* appeared to be epistatic towards to the *velB* (Fig 7A-7D). In sum, the application of salt stress to the corresponding *velB* and *msnA* deletion strain compared to wildtype revealed a positive implication of MsnA but not for VelB on fungal growth, independently of the fungal developmental program. Little is known about the regulation of salt-stress gene regulation by the MsnA in *A. nidulans*, whereas the binding profiling of Msn2 from *S. cerevisiae* (as ortholog of MsnA) during oxidative stress has been studied years ago [20,21]. *A. nidulans* MsnA might be able to occupy promoters of major players of the salt-stress signaling, even without any particular salt conditions. The ChIP-seq data generated for the current study were searched for in vivo binding events of MsnA to promoters of known salt-stress regulators. The promoters of *atfA* and *aslA,* encoding for the bZIP-type AtfA and the C2H2-type zinc finger AslA transcription factors correspondingly, both implicated in salt-stress response of *A. nidulans* [52,53], were found to be strongly occupied by the MsnA (Fig 7E and 7F). A subsequent gene expression analysis of the *atfA* and *aslA* in the Δ*msnA* and Wt strains from mycelia grown under vegetative conditions was performed. The results from this analysis, showed that *atfA* and *aslA* expression was strongly increased in the Δ*msnA* compared to wildtype strain (Fig 7G and 7H). These results further confirmed that those in vivo regulatory interactions of MsnA with the promoter of *atfA* and *aslA*, have an actual biological significance. All those findings together propose that in the absence of salt-stress, MsnA even prevents the unreasonable initiation of a salt-stress response by attenuating the expression of *atfA* and *aslA* genes encoding for salt-response regulators.

**Fig 7.**
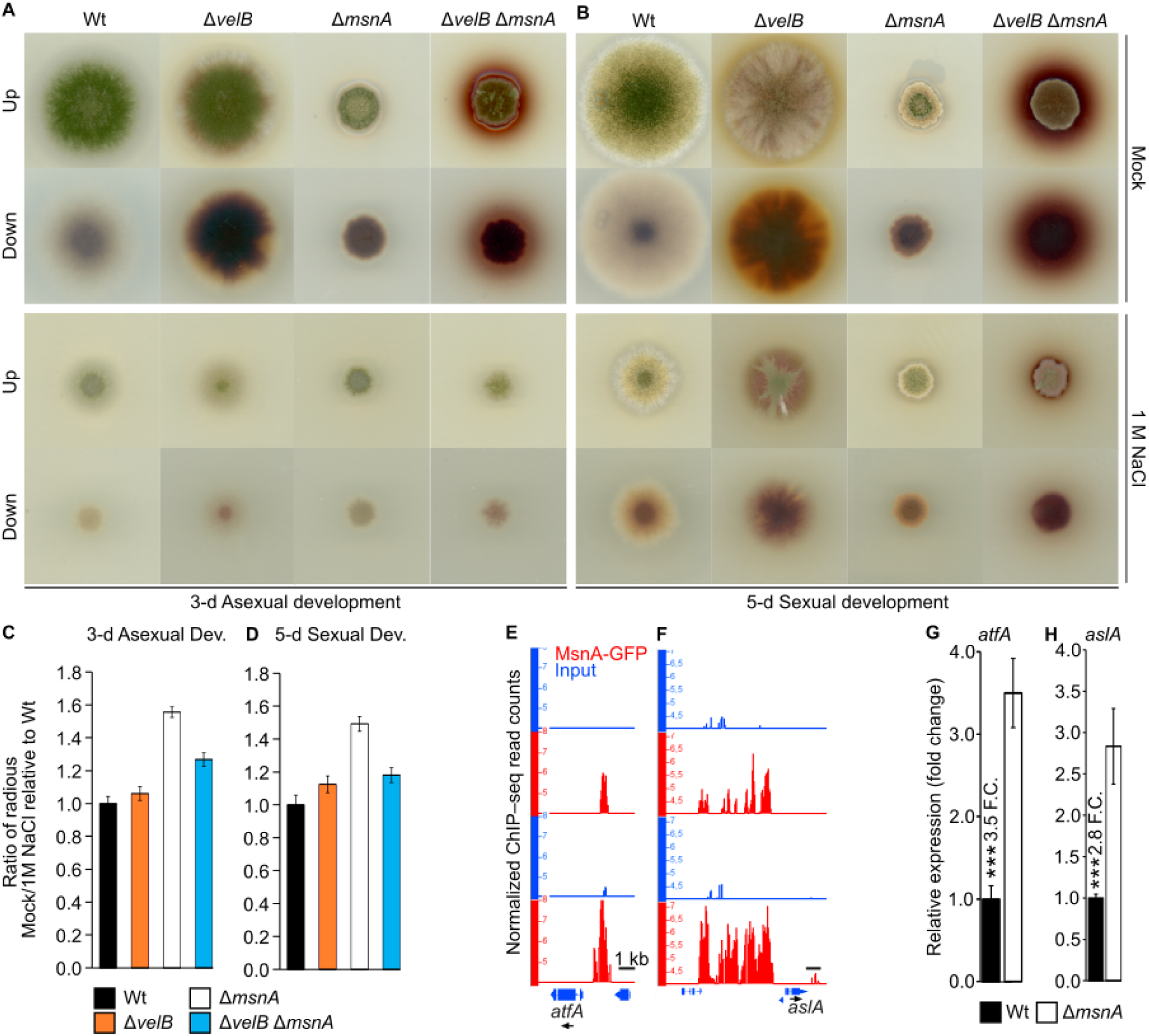
MsnA, but not VelB, mediates salt-stress response in *A. nidulans*. Phenotypical characterization of the Δ*msnA*, Δ*velB* and their corresponding double mutants under salt-stress. Scans of different strains under mock and 1 M NaCl treatment accordingly for 3-d asexual (**A**), and 5-d sexual (**B**) development. Initially 2000 spores were spotted in the middle of each medium plate. Bar diagrams show the ratio of the diameters of the colonies for different strains without (Mock) and with (1 M NaCl) the application of the salt treatment during 3-d asexual (**C**) and 5-d sexual (**D**) development. Presented data are averages and standard deviations of four biological replicates per each strain (under mock and treated condition. The screen shots from the Integrative Genome Browser (IGB) (**E** and **F**) shows a peak from the MsnA-GFP ChIP-seq data, where MsnA is associated in vivo with promoters of *atfA* and *aslA*, encoding for known regulators of the salt stress in *A. nidulans*. Gene expression analyses of *atfA* and *aslA* (**G** and **H**), under a regular vegetative growth (without salt-stress treatment), via qRT-PCRs, showing the strong repressive role of MsnA to the *atfA* and *aslA* gene expression. Presented gene expression data are averages and standard deviations of at least three biological replicates with each deriving by at least three technical replicates; Statistics made by Student’s t test: ***P ≤ 0.01.

### MsnA transcriptionally influences, directly and indirectly, methyltransferases with major roles in *A. nidulans* development

Sexual development is promoted by the trimeric complex of two velvet proteins, VeA and VelB, with the methyltransferase LaeA [4]. Asexual development is favored by the restriction of the VeA (a major sexual regulator) from the nucleus. The VapA-VipC-VapB complex is tethered to the plasma membrane via VapA under the dark, which favors the sexual growth. Under light, the VipC-VapB methyltransferase complex is released, and it restricts the entry of VeA in the nucleus by its physical interaction with it. VipC-VapB further enters the nucleus where then is affecting epigenetically the methylation status of histones, which leads to changes in the expression of genes that promote asexual development [6]. MsnA plays a major role in the asexual development, alongside with its molecular interplay with the velvet domain proteins. It was examined whether MsnA has a direct role in the regulation of those methyltransferases, which control either the asexual or the sexual developmental program of *A. nidulans*. Therefore, the MsnA ChIP-seq data were analysed for peaks nearby the genes of the methyltransferases. Sharp peaks where detected proximal to the gene of the Zinc finger VapA membrane protein, a subunit of the VapA-VipC-VapB complex and the VipC methyltransferase (Fig 8A). No peaks were detected for the other two methytransferases, VapB and LaeA. Next, we wanted to verify if those in vivo binding events of MsnA to the promoters of *vapA* and *vipC* are able to influence the expression of those genes. A gene expression analysis was performed with wildtype and Δ*msnA* strain. Vegetative growth and induction of fungal development on solid medium plates with or without light were analysed. The expression of *vapA* and *vipC* was increased in the Δ*msnA* strain versus the wildtype (Fig 8B), validating the binding of MsnA to the promoters of these genes (Fig 8A). When the expression of *vapA* and *vipC* was followed, at early stages of the asexual development, a different pattern emerged, particularly for the *vipC* expression. In two out of three time points tested, *vapA* was increased in the Δ*msnA* (Fig 8C), similar as was observed during vegetative growth (Fig 8B). In contrast, *vipC* was found to be repressed by MsnA during vegetative growth, whereas it was induced by MsnA under asexual growth conditions in all three distinct time points tested (Fig 8C).

**Fig 8.**
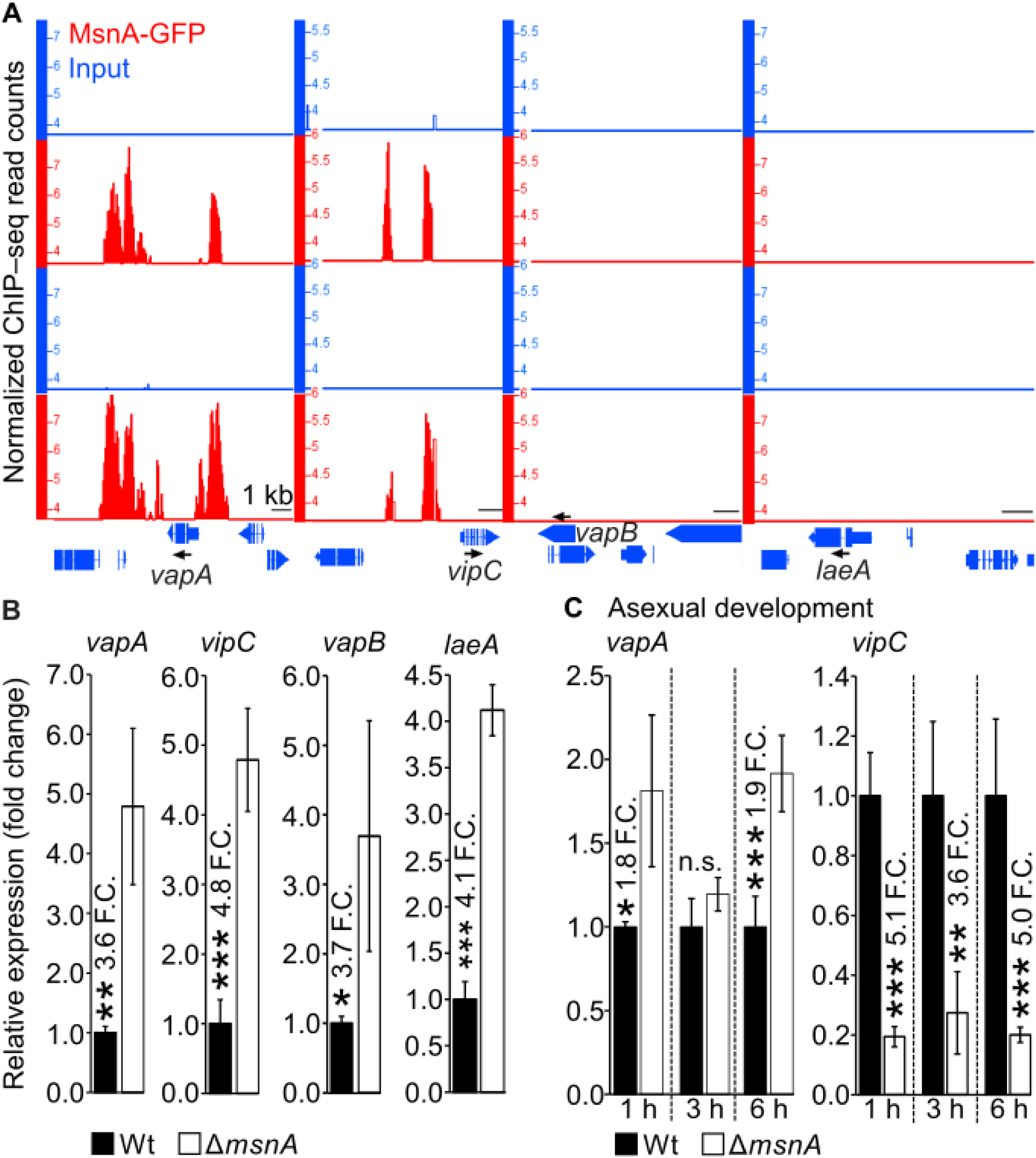
MsnA is directly in vivo to the promoters of *vipC* and *vapA*, encoding for parts of the VapA-VipC-VapB (asexual development promoting) methyltransferase complex. Screen shots (**A**) from the Integrative Genome Browser (IGB) depicting the presence or absence of MsnA-GFP from *vapA, vipC*, *vapB* and *laeA* promoter regions. Gene expression profiles (**B**) by qRT-PCR, for all the genes depicted in (**A**). All RNAs derived from mycelia of wildtype or Δ*msnA* strain, growing vegetatively in submerged cultures for 20 hours at 37° C shaking on a rotary shaker. Expression profiling (**C**) of *vapA* and *vipC* in mycelia derived from wildtype (Wt) and Δ*msnA* strains growing initially for 20 hours vegetatively and subsequently transerefed to solid medium plates under conditions favoring the asexual growth for the indicative time points. All gene expression data presented are average values and standard deviations of at least three biological replicates with each deriving by at least three technical replicates; Statistics made by Student’s t test: *P ≤ 0.05, **P ≤ 0.01 and ***P ≤ 0.01, n.s.: not significant.

Although MsnA has no preference to the promoters of the other two methytrasferases, VapB and LaeA, it was examined if it can influence the expression of their genes indirectly. Expression analysis of the *vapB* and *laeA*, for mycelia under vegetative growth, showed that indeed both genes were strongly repressed by MsnA (Fig 8B). In sum, MsnA has a new and previously elusive role as a transcriptional regulator of complexes between the velvet proteins and methyltransferases, with essential and established roles during the two distinct developmental programs of the fungus.

## Discussion

During its life span, *A. nidulans* has to respond to a variety of internal and external stimuli to coordinate and promote its development, that allows the production of spores, which secure its distribution and survival. Several characterized, but also plenty of yet uncharacterized TFs play major roles in this essential orchestration. Currently there are several examples where the role of a TF in various processes has been elucidated but their direct implication (for example, which are the direct target genes genome-wide) during the development remains elusive. *A. nidulans* MsnA falls into that category of TFs. As a C2H2-type zinc-finger TF, MsnA, has been linked to several stress responses such as those to salt/osmotic, temperature and hydrogen peroxide/oxidative (H_2_O_2_) stress [8,9]. There is lack of knowledge regarding the roles of MsnA during fungal development, specifically in a genome-wide scale. Here we show that MsnA, plays a major role in the development of *A. nidulans* under non-stress conditions: i) MsnA directly binds to promoters of genes encoding for master regulators in vivo, ii) MsnA fine tunes secondary metabolism and iii) MsnA interacts with other master regulators such as the velvet proteins and the VapA-VipC-VapB methyltransferase complex at the molecular level.

The evidence provided by the current literature, supporting a direct implication of MsnA in the development of *A. nidulans* under non-stress conditions are quite restricted. The facts supporting this hypothesis, show that the *msnA* deletion strain displays a severe restriction of the colony when grown in light. Nevertheless, the ability of this strain to form developmental structures and a quantification of those, was not further investigated [54]. Our data show that the deletion of *msnA* causes a very strong reduction in colony size, the numbers of asexual spores (conidia) and the number of the sexual structures (cleistothecia) that carry the sexual spores (ascospores) (Fig 1B-1C). Nevertheless, apart from the developmental delay that the Δ*msnA* shows compared to wildtype, it has to be mentioned here that both developmental programs remain functional and can be completed in the deletion strain (S1A and S1B Fig).

In yeast the preference of Msn2 to be associated with certain regions of the yeast genome under stress has already been studied the last decade. It has been shown that there is a specific dynamic interplay among Msn2 binding and the nucleosome occupancy that is crucial for the activation or (and) the repression of specific genes nearby [20,21]. However, in *A. nidulans*, up to now, not only an in vivo occupancy preference of MsnA (genome-wide) does not exist, but also the overall concept that MsnA, can operate as a TF, is up to now rather not clear. Even though it has been postulated that MsnA can control the expression of genes, there is hardly any data in literature where *A. nidulans* MsnA is directly associated with a promoter of a gene. MsnA can interact in vitro with the *mstB* promoter, presumably involved in transportation of monosaccharides that are necessary during sexual development [55]. In another filamentous fungus, *A. parasiticus*, MsnA is associated to specific promoters of genes involved in the synthesis of aflatoxin and stress response by in vitro binding assays [10]. Here we show that the MsnA is a nuclear localized protein, able to interact in vivo with many promoters of genes, across the genome of *A. nidulans*. Some of these genes encode major regulators of asexual or sexual development (Fig 3). In almost all of these cases, MsnA was binding directly in vivo in close proximity to the TSS of these genes. However, there were two striking examples where MsnA was associated with far distal regions from the TSS *brlA* and *sclB* as well (S3C and S3F Fig); although in the case of *sclB* there was also another clear peak in close proximity of its TSS. Because the expression of these genes was found to be severely affected in the Δ*msnA* strain compared to Wt, it can be postulated that their differential expression can either be due to indirect regulation or also because MsnA binds to those distal regions as well. Indeed, this remote binding in vivo affects their expression correspondingly. Nevertheless, a further in-depth examination is a subject of future studies. *A. nidulans* MsnA shares a similar DNA recognition motif with yeast Msn1 [20], so called STRE (stress response element). MsnA is located to the promoters of genes via its direct molecular interaction with these elements. As a result, their expression is influenced immediately (Fig 3). In sum, our findings highlight the role of MsnA as an important regulator for *A. nidulans* development, which can influence the expression of several known development-related genes directly, hence promoting both developmental programs.

Data, where orthologs of MsnA from other filamentous fungi (*A. parasiticus* and *A. flavus*) showed an association with secondary metabolism was related to the production aflatoxins and kojic acid [9]. Our broader approach elucidated that MsnA is directly or indirectly implicated, in the synthesis of secondary metabolites that are not only related to defense mechanisms of the fungus (such as DHMBA) but also to the regulation of its development (such as austinol and dehydroaustinol) (Fig 4). Therefore, these data underline a novel aspect of MsnA, acting as a major hub where stress signaling, secondary metabolism and development can be converged and coordinated, to help the fungus coping with various environmental stimuli. The coordination of developmental processes is rarely an act of a single player. Instead, it often is manifested as a collaborative work of several key regulators, which can collectively orchestrate the necessary actions, required for progression of the fungal development based on external and internal stimuli. According to this concept, it was discovered that MsnA is collaborating with other master regulators, to progress fungal growth. Working towards these lines a regulatory circuit among MsnA and velvet master regulators was disclosed. All velvet proteins were able to affect *msnA* expression. However, only *velB* and *vosA* expression were affected by MsnA. It was further shown that MsnA and VelB can mutually control each other’s expression, by in vivo association with their promoters (Fig 5). This molecular regulatory circuit among MsnA and VelB drove us to study the genetic interplay between the two genes. We discovered that *msnA* is epistatic to *velB* when the morphology and the size of the colony was examined independently from the developmental program (Fig 6). In sum, the novel molecular regulatory circuit among MsnA and VelB is further enhanced and supported by additional genetic interactions among these to regulators, to further orchestrate the development. Additionally, it was shown that an autoregulatory transcriptional loop might make MsnA to control its own expression (Fig 5A).

The VapA-VipC-VapB methytrasferase complex, is known for mediating the signal that induces asexual development in light from the plasma membrane into the nucleus [6]. This mediation involves the release of the VipC-VapB heterodimer from the plasma membrane, where is tethered via the VapA and its subsequent interaction with VeA. This prohibition of VeA, keeps it out of the nucleus, hence, makes it unable to interact with the nuclear LaeA methytransferase to initiate the sexual developmental program under light [6]. The reciprocal regulatory interplay between MsnA and the velvet proteins, alongside with the relationship that the VipC subunit of the trimeric methyltransferase complex shows with VeA [6], led to the hypothesis that there might be a link between MsnA and VapA-VipC-VapB as well. Our in vivo binding experiments showed that MsnA had no binding preference to the promoters of *vapB* and *laeA* but it was found to be strongly associated with the promoters of *vapA* and *vipC* (Fig 8A). These particular DNA-protein interactions found to influence the expression of the *vapA* and *vipC* genes directly and indirectly the *vapB* and *laeA* in a repressive manner during the vegetative growth. Based to the current model the interplay between the VipA-VipC-VapB and VeA-VelB-LaeA [4,6] complexes is balancing developmental programs of the fungus. Our findings enhance this model, by revealing another layer of transcriptional regulation, governed by MsnA, towards components from both of those complexes.

The role of MsnA in the stress signaling across many different fungal species is relatively well-established. Various stress stimuli can trigger the expression of *msnA*, or its paralogs from other fungi, among them salt/osmotic, oxidative and heat stress [8,9,56]. The discovery of the molecular and genetic network among MsnA and VelB, alongside the lack of knowledge on the potential implication of VelB in salt signaling in particular, formed a new hypothesis. In *A. nidulans* VelB together with MsnA might be able to mediate a salt-stress response. However, our data indicate that VelB does not contribute to a salt-stress response. Moreover, strains that lack *msnA* show a strong retardation in colony formation compared to wildtype during growth with 1 M NaCl. The study of the double mutant of *msnA* and *velB* did not reveal any significant additive or synergistic effect of the two transcription factors during salt-stress response, independently from the developmental program. Nevertheless, it did show that *msnA* is epistatic to *velB* (Fig 7A-7D). However, data from *A. niger* showed that the *msnA* deletion strain grown with 1 M NaCl had no significant effect on the formation of the colony compared to wildtype [57], suggesting perhaps that the important role of MsnA against salt stress in *A. nidulans* is not conserved in *A. niger*.

Our study revealed another novel role of MsnA related to the control of stress response. In particular, it was found that genes encoding for major stress regulators, such as AtfA [52,58,59] and AslA [53], can be repressed by MsnA in vivo, in the absence of any stress stimuli (Fig 7C-7H). Taken the results together, it becomes clear that the role of MsnA is not solely restricted to the mediation and orchestration of salt-stress response, but also to the deactivation of the corresponding signaling, by attenuating the expression of stress master regulators, when stress stimuli are no longer present. Based on these findings, it would be interesting to examine in a genome-wide scale, the in vivo binding preferences of MsnA under salt-stress or other stress conditions to see which are the genes, directly controlled by MsnA.

We propose a model (Fig 9) where MsnA functions as an organizer of a transcriptional hub. Known master regulators of development and stress responses, such as the velvet proteins, but also other so far unknown regulators, can be influenced by MsnA or (and) can affect MsnA as well. Through these mutual transcriptional interactions at the MsnA regulatory hub, the orchestration of *A. nidulans* development, secondary metabolism and stress response can be coordinated accordingly based on signals that the fungus receives by its environment.

**Fig 9.**
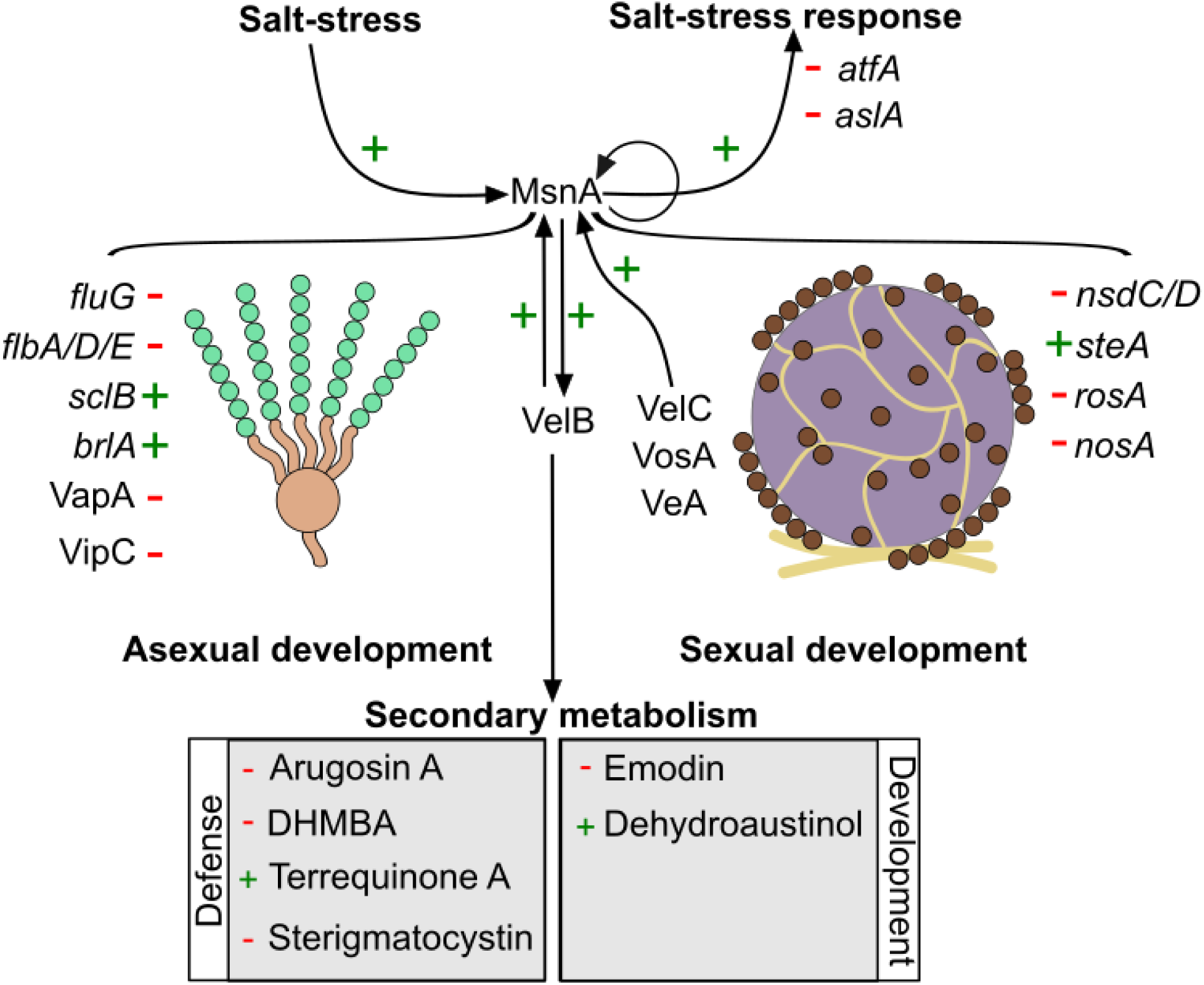
MsnA controls master regulators as well as regulatory gene networks to mediate developmental and salt-stress signals in *A. nidulans*. The illustrated model shows the global influence that MsnA has in different facets of *A. nidulans*′ biology. Master regulators of the asexual (*fluG*, *flbs*, *sclB*, *blrA*, *vapA* and *VipC*) as well as sexual (*nsdC*/D, *steA*, *rosA* and *nosA*) development, are directly (in vivo) transcriptionally controlled by MsnA, starting already from the vegetative growth of the fungus. A dynamic, mutual cross-regulatory circuit between MsnA and VelB coordinates and mediates several signals into organized transcriptional responses. Although, stress responses (such as high concentration of NaCl) are solely mediated by MsnA without any implication of VelB. MsnA seems to keeps the salt-stress signalling off, when no real stimulus is present, by the direct repression of crucial stress TFs such as AtfA and AslA. Secondary metabolism is also influenced by the function of MsnA. Transcriptional changes, provoked by MsnA (directly or indirectly), eventually lead to changes in of the final secondary metabolites (related to the defense or the development) of the corresponding metabolic pathways. Green crosses (+) depict the induction and red hyphens (-) the repression, caused by the direct in vivo association of MsnA to promoters of the corresponding genes during vegetative growth.

## Materials and methods

### Strains, media and growth conditions

The strains of *Aspergillus nidulans* and *Escherichia coli* used in this study are listed in the S1 Table and S2 Table correspondingly.

The *A. nidulans* strain that was used as a wildtype for all the experiments was AGB551. *A. nidulans* strains were cultivated in minimal medium with the composition of 1% (w/v) glucose, 2 mM MgSO_4_, 1x AspA (7 mM KCl, 70 mM NaNO_3_, 11.2 mM KH_2_PO_4_, pH 5.5), 0.1% (v/v) trace element solution (76 µM ZnSO_4_, 178 µM H_3_BO_4_, 25 µM MnCl_2_, 18 µM FeSO_4_, 7.1 µM CoCl_2_, 6.4 µM CuSO_4_, 6.2 µM Na_2_MoO_4_, 174 µM EDTA) and pH 5.5 [60]. For liquid medium there was a supplement of 0.1% (v/v) pyridoxine and 5 mM uridine. For solid medium 2 % agar and 0.1% (w/v) uracil was added. After the transformation, *A. nidulans* protoplasts were plated upon solid medium additionally containing 120 mg/mL nourseothricin as a selection marker. The selection-marker cassette was recycled from the A. *nidulans* genome after the selection process was done, by growing the selected strains on medium plates containing 0.5% (w/v) glucose and 0.5% (w/v) xylose.

The DH5α strain of *Escherichia coli* was used for cloning purposes of this study. The culture medium used for then cultivation of *E. coli* was a lysogeny broth (LB) [61] medium (1 % tryptone, 0.5 % yeast extract, 1 % NaCl, supplemented with ampicillin (100 µg/mL as a final concentration) for the selection of transformants with the desired plasmid. In case of solid medium an addition of 2 % (w/v) agar was used as well.

### Plasmid construction, handling and preparation

The extraction of the plasmid DNA was performed by the NucleoSpin® Plasmid Kit (Macherey-Nagel, Düren, Germany), under the manufacturer’s instructions. The final plasmids, carrying the constructed cassettes were verified via Sanger sequencing by Microsynth Seqlab GmbH (Göttingen, Germany) and the use of the Lasergene software (DNA Star Inc., Madison, USA).

The backbone plasmid used for the construction of the cassettes, which were transformed into *A. nidulans* strains, was the pBluescript KS (pME4696), constructed and used initially by [62]. This plasmid carries a *Pml*I restriction site for the integration of the target genes 5′-flanking region and a *Swa*I site for the integration of its 3′-flanking region. The 5′ and 3′ flanking regions define the exact position in the *A. nidulans* genome, where the incorporation of the constructed cassette will take place via homologue recombination, after transformation into *A. nidulans*. For the amplification of the 5′and 3′flanking regions of all the cassettes used in this study, genomic DNA from the AGB551 wildtype strain was used as a template. To construct the final cassettes, both flanking regions were inserted into the pME4696 vector via the GeneArt Seamless Cloning and Assembly Kit (Invitrogen, Carlsbad, CA, USA), following the manufacturer’s instructions. The cassettes included a recyclable selection marker, which provided resistance to nourseothricin (natRM).

### Construction of the deletion cassette of *msnA* and the *A. nidulans* Δ*msnA* strain

The amplification of the 5’-flanking region was performed with the MB62/63 pair of primers, yielding a 1663 bp amplicon, spanning from just before an ATG triplet, encoding for the TSS (translation start site) of the *msnA* and extended upstream from that point. The amplification of the 3’-flanking region of the deletion cassette was performed with the MB64/65, which gave an amplicon of 1036 bp starting just after a TGA triplet encoding for the stop codon of *msnA* and extending downstream from that point. Next, the 5’ region and 3’ region flanking regions were cloned into the pME4696 empty vector, giving the final plasmid of pME5558. Final constructs carrying both of the flanking regions were verified by Sanger sequencing, prior to the excision of the cassette with the *Mss*I (*Pme*l) restriction enzyme from the pMB4-10 plasmid, that was used subsequently for transformation into the *A. nidulans* wildtype strain, resulting in the AGB1707 strain, after the selection marker was recycled.

### Construction of the complementation cassette *msnA:hinge:GFP:3*’*UTR_msnA_:trpC-terminator* and the *A. nidulans* Δ*msnA/*MsnA-GFP strain

The first fragment of this cassette is composed by several different parts amplified from different templates fused by fusion PCR based on fragment compatibility introduced during the primers design. In short, the initial amplicon-A including part of the promoter region of the *msnA* and the *msnA* gene was amplified using genomic DNA from AGB551 as a template and the primer pair of MB1029/1030, yielding a 5032 bp fragment (the triplet encoding for the stop codon of *msnA* was removed via primer designing). Next, using as a template plasmid DNA, derived from the vector pChS459, the hinge-GFP fragment was amplified with the primers MB1031/1032 giving a 752 bp amplicon-B; hinge is a short (15 bp) spacer/non-coding part that simply distances the GFP from the protein that is going to be tagged with. At this point, amplicon-A and -B were fused by fusion PCR using the primer pair of MB1029/1032 yielding amplicon-C with the size of 5784 bp. Subsequently, an amplicon was generated, from genomic DNA of AGB551, with the primers MB1033/1034 in order to amplify specifically the 3′-UTR (as it was predicted in FungiDB databse) of *msnA* producing amplicon-D. The amplification of a *trpC*-terminator by using plasmid DNA from pChS459 as a template and the primers MB1035/1036 followed, yielding a 1460 bp amplicon-E. In the next step the amplicons-D and -E were fused by using the primers MB1033/1036 giving a product of 1460 bp, amplicon-F. In the last step of the construction of the first part for this cassette, the amplicons -C and -F were fused by fusion PCR by using the primer of MB1029/1036 yielding a final fragment of 7244 bp. The 3′-flanking region of the gene, was amplified from genomic DNA with the primers MB1037/1038 and had a size of 2001 bp. The first fragment and the 3’ region flanking region were finally cloned into the pME4696 empty vector leading to the pME5559 vector. Final cassettes were verified ed by Sanger sequencing, prior its excision with the *Mss*I (*Pme*l) restriction enzyme. The derived cassette then was subsequently transformed into the *A. nidulans* AGB1707 (Δ*msnA*) strain, resulting in AGB1708, after the selection marker was recycled.

### Construction of the *A. nidulans* Δ*msnA* Δ*velB* double deletion strain

The cassette that was used for the construction of the Δ*msnA* (as described previously), was also used for the construction of the double deletion strain. The corresponding cassette was excised, with the *Mss*I restriction enzyme from the pME5558 plasmid. It was then transformed and integrated in locus into the genome of the Δ*velB* (AGB1064) strain (sharing the same genetic background with the AGB551 wild type the except the deletion of the *velB*). Positive clones from the corresponding transformation were selected, subsequently the selection marker was recycled and the strains checked genetically for carrying both deletions (Δ*msnA* and Δ*velB*) by Southern hybridization, leading finally to the AGB1709 *A. nidulans* strain.

### Phenotypical characterization

To examine the development of the fungus, 2000 for each strain were point inoculated in the middle of a MM plates, supplemented 0.1% (v/v) pyridoxine, 5 mM uridine, and 0.1% (w/v) uracil. After growth under asexual or sexual conditions for the indicated time, scans were made from the top and bottom of the plates.

For single cleistothecia images and close shots of the colonies it the Axiolab microscope (Carl Zeiss Microscopy, Oberkochen, Germany) and the SZX12-ILLB2-200 binocular microscope (Olympus, Shinjuku, Japan) were used.

For quantification of conidia, identical number of spores from all strains were equally distributed incubated on solid medium and incubated for the indicated time under asexual growth conditions. The spores from each strain were collected and measured with the Coulter Z2 particle counter (Beckman Coulter GmbH, Krefeld, Germany).

For the quantification of cleistohecia, identical number spores from all strains were equally distributed in supplemented (0.1% (v/v) pyridoxine, 5 mM uridine, and 0.1% (w/v) uracil) MM plates and grown for indicated time under sexual growth conditions. Subsequently three agar plugs of 5 mm^2^ were removed from each plate (by using the larger side of 200 μL pipette tip) and placed on a new agar plate. The exact number of cleistothecia for each strain was then assessed by counting.

### Chromatin immunoprecipitation, Sequencing and data analysis

#### Chromatin immunoprecipitation (ChIP)

The Chromatin immunoprecipitation (ChIP) experiments were performed with mycelia derived from submerged cultures. In the case of the ChIP-seq for the MsnA, the *ΔmsnA* MsnA-GFP complementation strain was used, expressing *msnA* fused with *GFP* (*Green Fluorescent Protein*), under its native promoter. In the case of the ChIP for the VelB, the VelB-GFP strain was used, expressing *velB* fused with *GFP*, under its native promoter and the Wt strain as a negative control. In both cases, a total number of 5×10^8^ spores were inoculated 500 mL liquid medium inside 2 L flasks and grown for 20 hours under constant rotation, light and at 37 °C. Subsequently, mycelia were harvested quickly, dried and immersed in fixation solution containing 1% formaldehyde for 20 minutes. The whole ChIP experiments, were performed as described in Sasse *et al.,* 2023 [63], with minor modifications for each ChIP, which are the following. For the ChIP-seq of MsnA, two independent biological replicates of *ΔmsnA* MsnA-GFP were used. As inputs (the negative controls) the samples after shearing of the chromatin without GFP antibody added were considered. For the ChIP with the VelB, three independent biological replicates were used from the VelB-GFP strain and another three from the Wt strain (the negative controls); the IPs of both strains were performed with GFP antibody. For the ChIP with MsnA the input samples were subjected to the same DNA purification procedure as the samples derived from the IP with the GFP antibody, prior to library preparation for sequencing. For the VelB ChIP, a GFP antibody was applied to the IPs derived from VelB-GFP and Wt strains. All reagents, instruments, kits and rest of the procedures for both of the ChIP experiments were the same as in Sasse *et al*., 2023 [63].

#### Library preparation and NGS sequencing

ChIP-seq libraries preparation and the following sequencing were performed at the NGS-Integrative Genomics Core Unit (NIG), University Medical Center Göttingen. Initially, the quantity and quality of ChIPed-DNA and input samples were determined by a Fragment Analyzer. The preparations of the ChIP-seq libraries was done by using the TruSeq ChIP Library Preparation Kit (Illumina, San Diego, USA), following manufactures′ instructions. The size range of the final DNA libraries were assessed with the Fragment Analyzer, using the SS NGS Fragment 1-6000 bp Kit, with an average size of 340 bp. The Denovix system (Bio-Rad Laboratories, Hercules, CA, USA) was used for the quantification of DNA libraries. Libraries were amplified and sequenced on an S1 flow cell NovaSeq 6000 (Illumina, San Diego, USA), for 100 cycles. The produced sequencing images were processed with the BaseCaller Illumina software to generate BCL files. Those files were then demultiplexed into fastq files with bcl2fastq v2.20.0.422 generating a FastQC for data quality control.

The subsequent ChIP-seq analysis was performed partly by using the same software and pipelines as presented in Sasse *et al*., 2023 [63]with the minor modifications that are following. Part of the analysis was performed in the GALAXY platform [64] maintained by the GWDG (Gesellschaft für wissenschaftliche Datenverarbeitung mbH Göttingen). For the mapping of the raw sequences, derived from the high-throughput sequencing, the *Aspergillus nidulans* genome (downloaded from fungidb.org: FungiDB-46_AnidulansFGSCA4_Genome.fasta) was used. The raw sequencing data for the ChIP-seq experiment have been deposited at NCBI [BioProject ID PRJNA1198984].

### DNA extraction

The genomic DNA derived from *A. nidulans* mycelia grown overnight in liquid cultures under constant rotation, at 37 °C and in light. The extraction of genomic DNA was performed based to the protocol described by Thieme *et al*., 2018 [44]. All the concentrations of the purified DNAs were measured by NanoDrop ND-1000 spectrophotometer (Peqlab, Erlangen, Germany).

### Transformations *E. coli* and *A. nidulans*

Trasnformation of *E. coli* and *A. nidulans* strains were performed based on protocols described in Meister *et al*., 2019 [62]. Successful transformation of cassettes into the *A. nidulans* genome were confirmed by Southern hybridization as described by Southern, 1975 [65]. For Southern’s probe labelling the AlkPhos Direct Labeling Module (GE Healthcare Life Technologies, Little Chalfont, UK) was used following the manufacturer’s instruction.

### RNA extraction and cDNA synthesis

All strains were initially inoculated with the same number of spores (10^8^) in 100 mL liquid medium. After 20 hours of vegetative growth, mycelia were dried in miracloth (Merck KGaA, Darmstadt, Germany).100 mg were quickly weighted for each biological replicate and placed inside 2 mL reaction tubes with three zirconium oxide beads, SiLibeads (Sigmund Lindner GmbH, Warmensteinach, Germany), then were frozen instantly in liquid nitrogen. In case of the RNA that deriving from asexual and sexual conditions, the mycelia were transferred on solid MM after initial 20 hours of vegetative growth plates which were incubated for the indicated time under asexual (light, 37 °C) or sexual (dark, 37 °C, parafilm around the plates) conditions. For each time point, mycelia were harvested and handled as described before. Sampled mycelia were grinded for 1 minute with a ball mill MM400 (Retsch, Haan, Germany), just before RNA extraction; the teflon cassettes-cases (2 mL epis holders) of the mill were precooled in -80°C for about an hour prior the placement of the frozen 2 mL with the sample’s mycelia into them for grinding. RNA extraction followed with the pulverized mycelia by using the RNeasy Plant Miniprep Kit (Qiagen) according to manufacturer’s instructions. The final concentration of the RNAs was assessed by using the NanoDrop ND-1000 spectrophotometer (Peqlab, Erlangen, Germany). The subsequent synthesis of cDNA was performed with 1.0 µg total RNA from each sample by using the QuantiTect Reverse Transcription Kit (Qiagen, Hilden, Germany) according to the manufacturer’s protocol.

### Quantitative real-time PCR

Quantitative real-time polymerase chain reaction (qRT-PCR) was employed for studying gene expression. The CFX Connect Real-Time System (Bio-Rad Laboratories, Hercules, CA, USA) was used. All cDNA templates were diluted 1:5 for the qRT-PCR, which was performed with the MESA GREEN qPCR MasterMix Plus for SYBR from EUROGENTEC (Lüttich, Belgium) by following the manufacturer’s instructions. Data was partially analysed with the CFX ManagerTM 3.1 software package from Bio-Rad Laboratories and partially with Excel (Microsoft, *Washington, USA*), using the ΔCt method with a reference gene 2Ct(reference)-Ct(target) (BioRad Laboratories, qRT Application Guide) method for relative quantification of gene expression. As reference gene, *h2A* (*histone2A*) was used. qRT-PCR experiments were conducted at least with a minimum of 3 biological replicates, each consisting of at least three technical replicates. All qRT-PCR primers, used in this study are listed in the S3 Table.

### Secondary metabolites analysis

A total number of 10^5^ spores were equally spread on solid medium. Colonies were grown for the indicated time under asexual and sexual growth conditions. For each strain, three plates were prepared (three replicates). The sampling, SM extraction, LC-MS and the subsequent analysis was performed as described by Liu *et al*, 2021 [46]. Specific details for all the all the detected secondary metabolites are listed in the S5 Table.

### Microscopy

For fluorescence microscopy, 3500 spores of each strain were inoculated in 400 µL liquid medium with supplements, placed in a single well of 8-well chambered coverslip (Ibidi GmbH, Gräfelfing, Germany), incubated at 37 °C and light for 20 hours. Fluorescence microscopy was performed with the inverted confocal microscope Zeiss AxioObserver, Z.1 (Zeiss, Oberkochen, Germany), and the software SlideBook 6.0 software package (Intelligent Imaging Innovations GmbH, Göttingen, Germany). For staining of nuclei the Hoechst dye (Invitrogen, Massachusetts, USA) was used.

### Figures processing

The processing of the all figures was done by the vector-graphics editor Inscape (Inkscape Project, 2020; Inkscape, available at https://inkscape.org).

## Acknowledgments

We thank Dr. Gabriela Salinas and Fabian Ludewig from the NGS-Integrative Genomics Core Unit (NIG), University Medical Center Göttingen for excellent support performing NGS-based approaches. We also thank Blagovesta Popova for her comments on our manuscript. We acknowledge support by the Open Access Publication Funds of the Göttingen University.

## Supporting information

**S1 Fig. The MsnA regulator is a cytoplasmatic and a nuclear localized protein, which strongly affects growth’s colony formation during the development of *A. nidulans*.**

**S2 Fig. Proteins associated with the cell periphery, the plasma membrane and the extracellular region, are enriched among the top targets of MsnA.**

**S3 Fig. MsnA influences the expression of genes encoding established *A. nidulans* master regulators in mycelia growing under asexual and sexual growth conditions.**

**S1 Table. *A. nidulans* strains used in this study.**

**S2 Table. *E. coli* used in this study.**

**S3 Table. Primers used in this study.**

**S4 Table. ChIP-seq common locus IDs with peaks up to 3 kb promoter regions.**

**S5 Table. Secondary metabolites controlled by MsnA under asexual growth conditions, as detected by detected by LC-MS/MS.**

